# Adipose Triglyceride Lipase is needed for homeostatic control of Sterol Element-Binding Protein-1c driven hepatic lipogenesis

**DOI:** 10.1101/2020.11.02.363440

**Authors:** Beatrix Irene Wieser, Paola Peña de la Sancha, Silvia Schauer, Helga Reicher, Wolfgang Sattler, Rolf Breinbauer, Martina Schweiger, Peter John Espenshade, Rudolf Zechner, Gerald Hoefler, Paul Willibald Vesely

## Abstract

Sterol Regulatory Element-Binding Protein-1c (SREBP-1c) is translated as an inactive precursor-protein that is proteolytically activated to promote fatty-acid (FA) biosynthesis, when unsaturated (u)FAs are scarce. During fasting, however, lipogenesis is low, and adipose-tissue lipolysis supplies the organism with FAs. Adipose TriGlyceride Lipase (ATGL) is the rate-limiting enzyme for lipolysis, and it preferentially hydrolyzes uFAs. Therefore, we hypothesized that ATGL-derived FAs may suppress the proteolytic activation of SREBP-1c in the liver. Here we show that (i) SREBP-1c is inactive during fasting but active after refeeding, (ii) uFA species liberated by ATGL suppress SREBP-1c activation *in vitro*, (iii) SREBP-1c is hyper-activated in livers of mice lacking ATGL, and (iv) pharmacological inhibition of ATGL selectively activates SREBP-1c in hepatocytes. Our findings highlight an ATGL/SREBP-1c axis, instrumental to coordinate lipogenesis and lipolysis, whose homeostatic regulation is crucial to avoid severe diseases including diabetes, cardiomyopathy, and even cancer.

## Introduction

Sterol Regulatory Element-Binding Protein-1c (SREBP-1c) is transcriptionally driven by carbohydrate rich diets and translated as an inactive membrane-bound precursor (P)-SREBP-1c. If unsaturated (u) fatty-acids (FAs) and/or sterols are scarce, P-SREBP-1c traffics to the Golgi apparatus (Golgi) together with its chaperone SREBP Cleavage-Activating Protein (SCAP). There, its N-terminal transcription-factor domain (N)-SREBP-1c is proteolytically released from the membrane. Subsequently, N-SREBP-1c migrates to the nucleus, where it drives the transcriptional program for lipogenesis (Hannah, Ou, Luong, Goldstein, & Brown, 2001; J.D. Horton, Bashmakov, Shimomura, & Shimano, 1998; J. D. Horton, Goldstein, & Brown, 2002; Matsuda et al., 2001). Consequently, low-fat/high-carbohydrate diets (HChD) induce FA and triglyceride (TG) synthesis. During fasting, however, lipogenesis is low and adipose-tissue lipolysis is strongly activated by lipolytic hormones and low plasma insulin levels (Zechner, 2015). This results in FA release from adipose-tissue TG stores, increases plasma FA concentrations, and elevates FA uptake by the liver. Adipose TriGlyceride Lipase (ATGL) catalyzes the first and rate-limiting step of TG-lipolysis, and importantly, it preferentially hydrolyzes uFAs from TG. ATGL-knockout animals, in turn, are lipolysis-defective and show only half the fasting plasma non-esterified FA (NEFA) levels of controls. The uFAs, palmitoleic acid (16:1), oleic acid (18:1) and linoleic acid (18:2), are even further underrepresented (Eichmann et al., 2012; Haemmerle et al., 2006). Notably, uFAs were found to suppress the proteolytic activation of P-SREBP-1c through stabilization of the ER anchor-protein of the SCAP/SREBP complex, Insulin-Induced Gene-1 protein (INSIG-1) (Lee et al., 2010; Lee, Song, DeBose-Boyd, & Ye, 2006; Lee, Zhang, Feramisco, Gong, & Ye, 2008). We hypothesized that adipose-tissue lipolysis-derived FAs may contribute to the regulation of SREBP-1c in the liver. To test this hypothesis, we analyzed the interplay between the master regulator of FA biosynthesis, SREBP-1c, and the rate limiting enzyme for lipolysis, ATGL, *in vitro* and *in vivo*.

## Results

During fasting, adipose-tissue lipolysis is strongly activated by lipolytic hormones and low plasma insulin levels (Zechner, 2015). To understand how the nutrient status affects SREBP-1c activation, we kept mice under normal chow diet, fasted them overnight, or fasted them overnight and re-fed them a high carbohydrate/low-fat diet (HChD), before sacrifice (Figure 1A). ER membrane resident P-SREBP-1c and nuclear localized N-SREBP-1c were detectable by western blot (WB) in livers of fed mice, reduced by fasting, and profoundly upregulated by refeeding (J.D. Horton et al., 1998; J. D. Horton et al., 2002). The two major SREBP-1c target genes, *Acetyl-CoA-Carboxylase-1* (*Acc-1)* and *Fatty Acid Synthase* (*Fasn)* reacted in line with N-SREBP-1c (Figure 1B). Thus, the lipogenic program in liver is suppressed by fasting and activated after HChD refeeding (J.D. Horton et al., 1998; J. D. Horton et al., 2002; Im et al., 2009). To investigate if uFA species liberated by ATGL during fasting (Eichmann et al., 2012) are able to suppress proteolytic activation of SREBP-1c, we generated a UF1c cell-line that constitutively expressed a flag-tagged SREBP-1c reporter protein (Hua, Sakai, Brown, & Goldstein, 1996). The uFAs (16:1, 18:1, 18:2, and 20:4) inhibited P-SREBP-1c processing in UF1c cells, while the sFA (16:0) showed no effect (Figure 1C). Hence, uFA species underrepresented in the fasting-plasma of ATGL knockout mice (Eichmann et al., 2012) suppressed SREBP-1c cleavage-activation, *in vitro* (Hannah et al., 2001).

**Figure 1:**
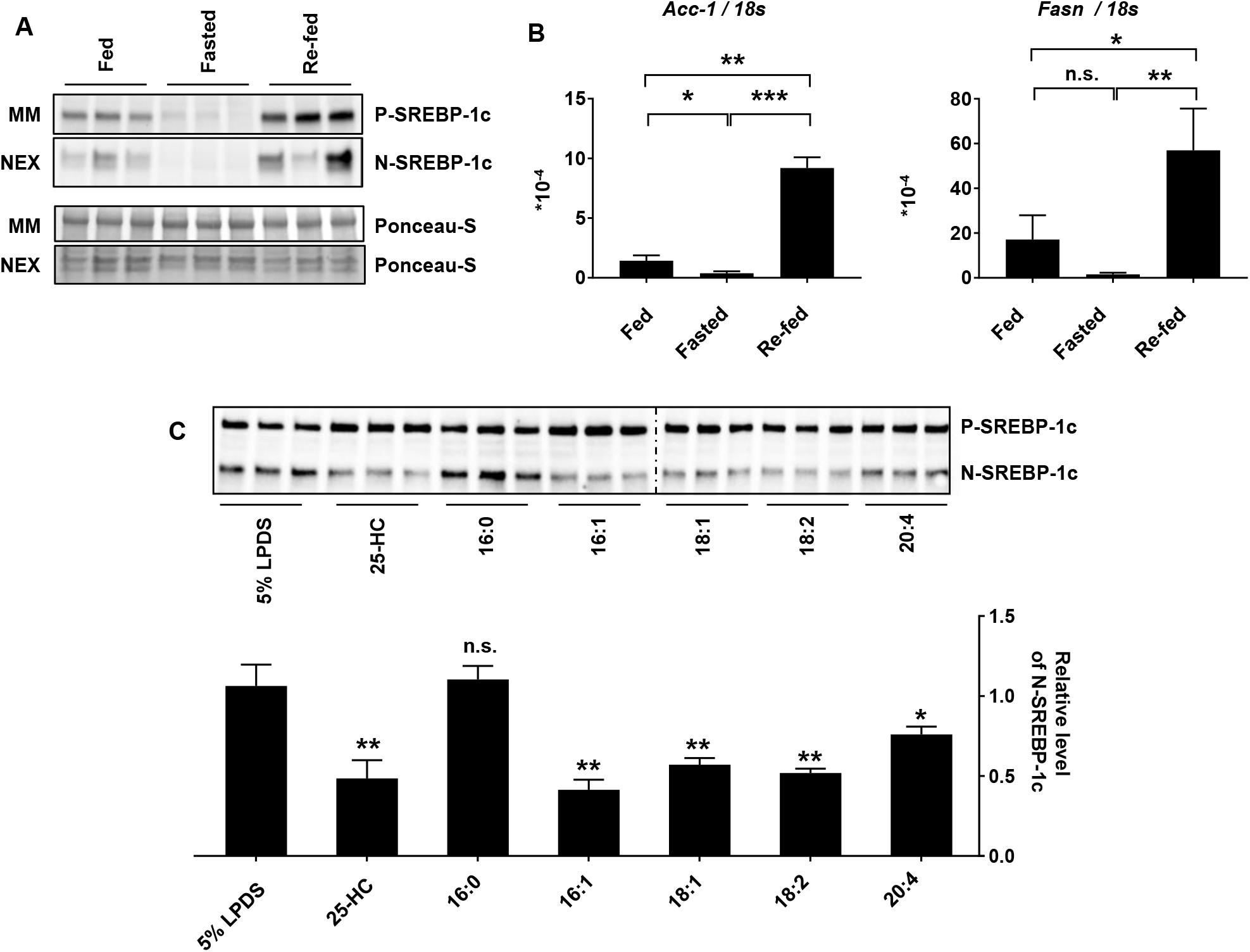
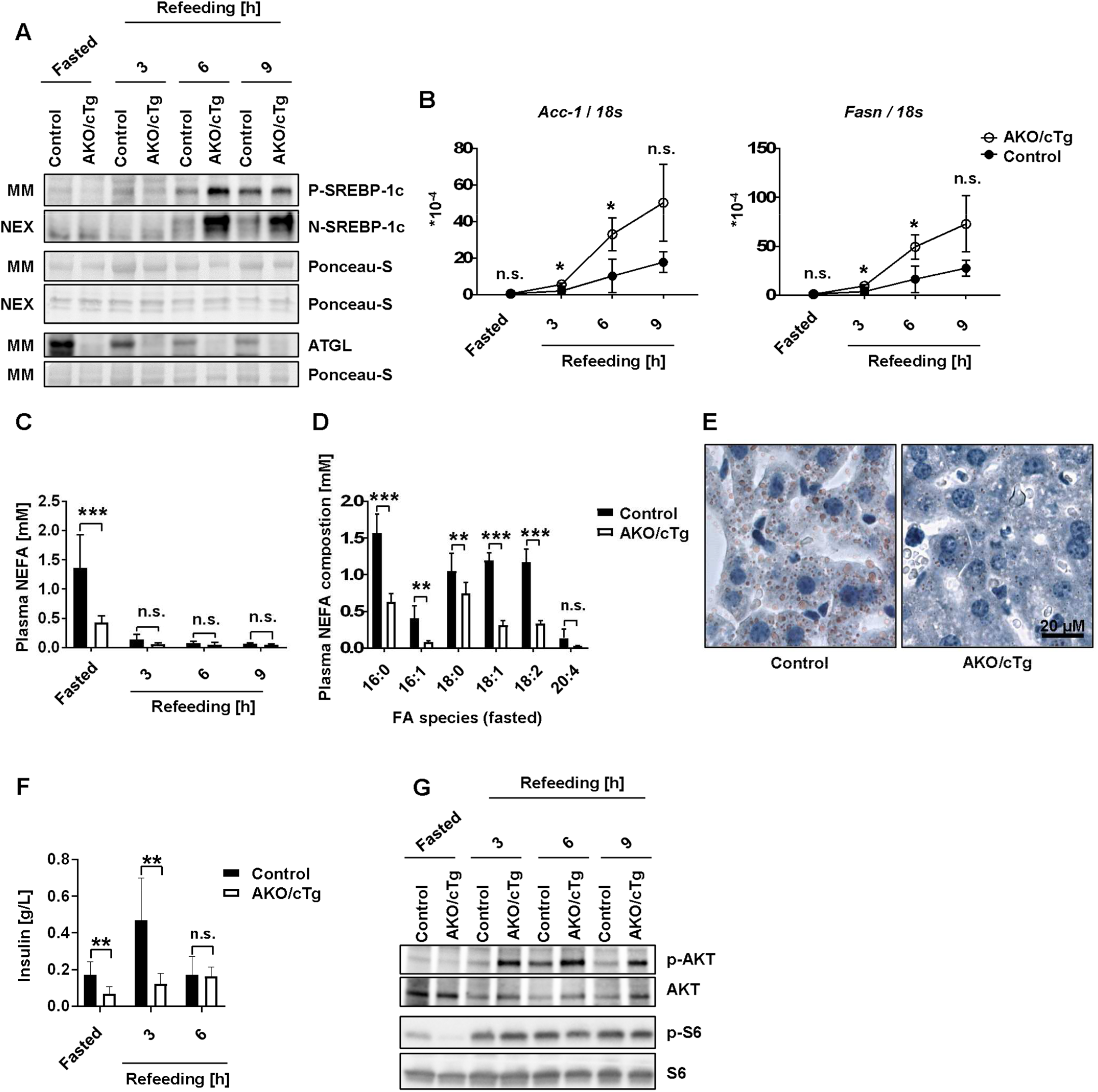
Regulation of SREBP-1c by feeding, fasting and fatty-acids (FA). **A)** Liver SREBP-1c precursor (P-SREBP-1c) and its transcriptionally active N-terminal (N) proteolytic fragment (N-SREBP-1c) are upregulated by food consumption. Mice were fed normal chow (Fed), fasted for 12 h (Fasted) or, fasted for 12 h and re-fed a high carbohydrate/low-fat diet (HChD) for 12 h (Re-fed). All mice were sacrificed together in the morning. Livers were resected, microsomal membrane fractions (MM) and soluble nuclear extracts (NEX) were prepared and subjected to western blot using SREBP-1c specific antibodies. Ponceau-S stained membranes are shown as loading controls. Each lane represents a liver from one mouse. **B)** Liver *Acc-1* and *Fasn* (mRNA) levels were determined by qPCR. n=3/group, technical replicates (tech rep) =2/sample. **C)** Unsaturated FAs decrease the relative level of N-SREBP-1c, compared to P-SREBP-1c, *in vitro*. We constructed a U2OS cell line constitutively expressing 2xFlag-SREBP-1c (UF1c). UF1c were seeded at 60% confluency using standard cell culture medium. 24 h later, the medium was replenished by 5% lipid depleted serum containing medium (5% LPDS) or 5% LPDS with 1 mg/ml 25-HC, or 100 µM FAs present, as depicted. Cells were incubated for 16 h and 2 h before harvest the protease inhibitor ALLN was added. Whole cell extracts were subjected to western blot. P- and N-SREBP-1c were detected, using anti-Flag antibodies. Band intensities were measured using ImageJ, NIH. Relative levels of proteolytically cleaved N-SREBP-1c were calculated as the relative fraction of N-SREBP-1c / P-SREBP-1c signal intensities, and are presented in the diagram below the western blot. n=3/group, tech rep=1/sample. Unpaired t-tests were used to compute significance levels, not significant, n.s.; p≤0.05 *; ≤0.01 **; ≤0.001 ***.

Our next aim was to test if uFA species liberated by ATGL during fasting (Eichmann et al., 2012) regulate SREBP-1c activation in mouse liver. To avoid the fatal heart phenotype of mice globally lacking *Atgl*, we used AKO/cTg, whole body *Atgl* knockout mice, with cardiac transgenic expression of *Atgl*, (*Atgl*^-/-,Myh6ATGL+/+^), and compared these to isogenic controls (*Atgl*^+/+,Myh6ATGL+/+^) (Haemmerle et al., 2011). In livers of fasted animals, no SREBP-1c specific signals were detectable by WB (Figure 2A). 6 h after HChD re-feeding, both N- and P-SREBP-1c signals were substantially stronger in the knockout group than in controls. P-SREBP-1c signal intensity returned to normal 9 h after refeeding, but N-SREBP-1c was still strongly over-activated in AKO/cTg compared to control livers. In line, the SREBP-1c target genes *Acc-1* and *Fasn*, were upregulated more rapidly and to a higher level in AKO/cTg compared to control livers (Figure 2B). As expected, ATGL was only detectable in control livers and steadily decreased after HChD refeeding (Figure 2A). As a result of lower FA availability in plasma (Figure 2C), largely due to a reduction of uFAs (Figure 2D), fasted AKO/cTg mice exhibited reduced Oil-red-O (ORO) neutral lipid staining in liver sections compared to controls (Figure 2E). Collectively, these findings suggest that uFAs liberated by ATGL may suppress SREBP-1c activation in the liver. However, insulin also triggers SREBP-1c activation (Matsuda et al., 2001; Owen et al., 2012), and the elevated insulin sensitivity of mice lacking ATGL (Haemmerle et al., 2006; Kienesberger et al., 2009; Schoiswohl et al., 2015; Schreiber et al., 2015a) could be an alternative explanation for increased SREBP-1c activation (Figure 2A). However, AKO/cTg mice showed little to no upregulation of their plasma insulin levels after re-feeding as opposed to isogenic controls (Figure 2F). Nevertheless, we could confirm elevated tissue specific insulin sensitivity in their livers, using a p-Ser473 AKT (p-AKT) specific WB antibody (Figure 2G, upper panels) (Kienesberger et al., 2009). SREBP-1c cleavage is, however, activated by insulin in a process requiring small ribosomal subunit protein S6 (S6) phosphorylation by its kinase, p70S6K (Owen et al., 2012). To test the p70S6K/S6 arm of insulin receptor signaling in the liver, we used a p-Ser440/44 S6 (p-S6) specific antibody. The p-S6 WB signal was downregulated in livers of fasted AKO/cTg mice compared to controls and yielded similarly strong signals after re-feeding (Figure 2G, lower panels). Hence, we conclude that the SREBP-1c over-activation in AKO/cTg mice is due to their lipolysis defect rather than due to their enhanced insulin sensitivity.

**Figure 2:**
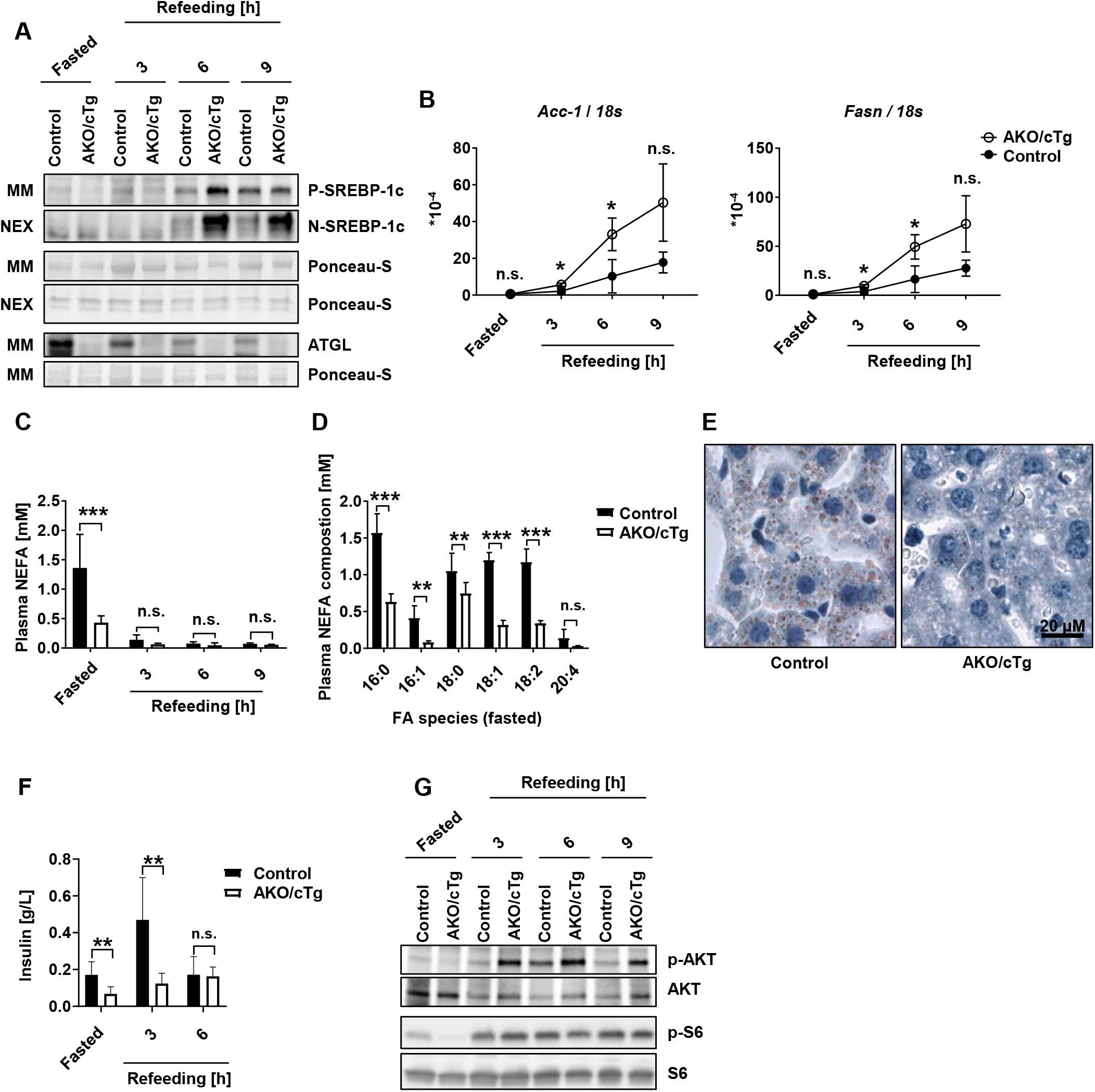
Regulation of SREBP-1c by systemic ablation of ATGL. *Atgl* knockout mice where the lethal cardiac phenotype was rescued by cardiac transgenic (cTg) ATGL expression (AKO/cTg) show considerably increased N-SREBP-1c levels, when compared to controls. Mice were fasted for 12 h overnight and subsequently groups of 3 mice were either sacrificed (Fasted) or refed an HChD and sacrificed at the time-points indicated (Refeeding). **A)** Livers were resected and microsomal membrane fractions (MM) and soluble nuclear extracts (NEX) were prepared and subjected to western blot. Respective extracts from 3 mice were pooled and analyzed using antibodies specific for the proteins indicated. Ponceau-S stained membranes are shown as loading controls. **B)** Liver *Acc-1* and *Fasn* (mRNA) levels were determined by qPCR. n=3/group, technical replicates (tech rep)=3/sample. **C)** Plasma non-esterified fatty-acid (NEFA) levels were determined biochemically. n≥3, tech rep=1. **D)** Plasma NEFA composition of fasted mice. n=5/group, tech rep=1-2/sample. **E)** Liver sections were prepared from 3 fasted mice per genotype and stained with Oil-red-O neutral lipid dye. Representative images are depicted. **F)** Plasma insulin concentrations were measured by Enzyme-Linked Immunosorbent Assay; n≥6/group, tech rep=1/sample. **G)** Liver MM fractions from (A) were analyzed by western blots using antibodies specific for the proteins indicated. Unpaired t-tests were used to compute significance levels, not significant, n.s.; p≤0.05 *; ≤0.01 **; ≤0.001 ***.

To directly test if adipose-tissue lipolysis derived FAs regulate HChD-induced SREBP-1c activation in the liver, we analyzed lipolysis defective, adipose-specific *Atgl*-deficient AAKO mice (*Atgl*^flox/flox,AdipoQCre+/-^) in comparison to control mice (*Atgl*^flox/flox^) (Schoiswohl et al., 2015) (Figure 3A). WB revealed weak P-SREBP-1c and no N-SREBP-1c signals in livers of fasted mice. After refeeding, P-SREBP-1c signal intensities gradually increased in control livers, but not in those of AAKO mice. The N-SREBP-1c signal increased in both genotypes with refeeding but after 6 h it was substantially stronger in AAKO livers than in those of control mice. In sharp contrast to mice globally lacking ATGL (described in Figure 2A), N-SREBP-1c levels were similar in both genotypes, at later timepoints. Accordingly, both SREBP-1c targets, *Acc-1* and *Fasn*, showed an expression peak 6 h after HChD refeeding in AAKO, but not in control livers (Figure 3B). ATGL was more abundant in AAKO livers compared to control, and steadily declined in both genotypes after refeeding (Figure 3A). Liver sections from fasted AAKO mice showed higher neutral lipid content than those of controls, as indicated by ORO staining and biochemical TG measurements (Figures 3C and D). Due to reduced adipose-tissue lipolysis, fasting plasma FA concentrations of AAKO mice were only half as high as in controls (Figure 3E) (Schoiswohl et al., 2015). As seen in the complete knockout model, GC/FID revealed a relative reduction of uFA in the fasting-plasma of the AAKO group compared to controls (Figure 3F). Tissue specific insulin signaling was slightly stronger activated in AAKO livers than in control livers 3 h post re-feeding, as indicated by p-AKT WB (Figure 3G, upper panels) (Kienesberger et al., 2009). However, the specific marker for insulin receptor signaling dependent SREBP-1c activation, p-S6 (Owen et al., 2012), was again weaker in the knockout group after fasting. During refeeding it showed no differences between the genotypes (Figure 3G, lower panels). Hence, our data indicate that adipose ATGL activity provides uFAs that temporarily suppress P-SREBP1c, for a few hours after HChD refeeding. Tissue-specific insulin signaling does not seem to cause this effect. Whether lipolysis-derived uFAs from adipose-tissue directly suppress SREBP-1c cleavage or, if uFAs are first esterified into TG and subsequently re-activated by liver-resident lipases, however, remains uncertain.

**Figure 3:**
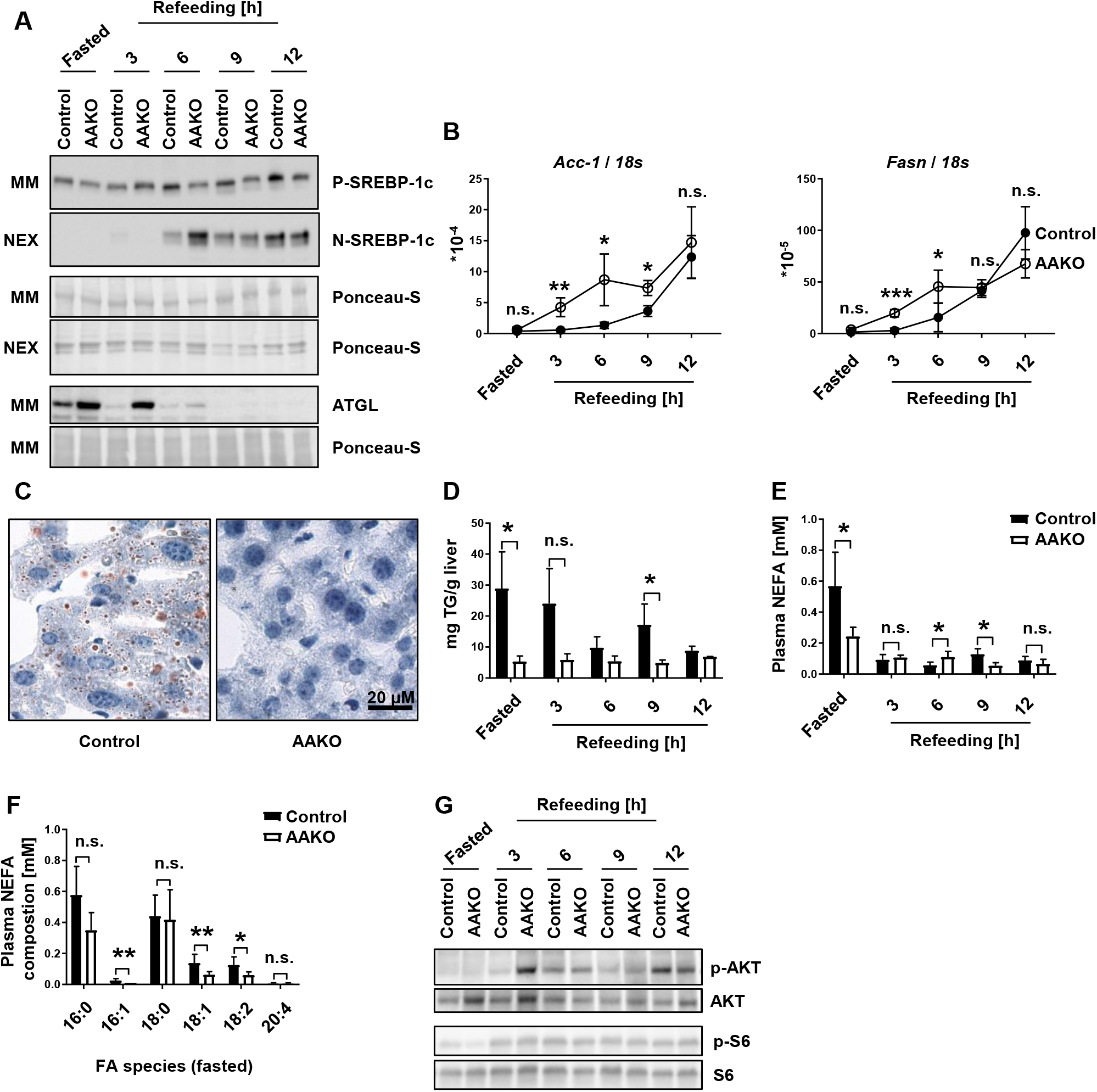
Adipose-tissue ATGL regulates SREBP-1c proteolytic-processing *in vivo*. Adipocyte specific knockout of *Atgl* (AAKO) leads to increased levels of N-SREBP-1c in livers during high carbohydrate/low-fat diet (HChD) re-feeding. Mice were fasted for 12 h overnight and subsequently groups of mice were either sacrificed (Fasted) or refed an HChD and sacrificed at the time-points indicated (Refeeding). **A)** Livers were resected and microsomal membrane fractions (MM) and soluble nuclear extracts (NEX) were prepared. Respective extracts from 3 mice were pooled and analyzed using antibodies specific for the proteins indicated by western blot. Ponceau-S stained membranes are shown as loading controls. **B)** Liver *Acc-1* and *Fasn* (mRNA) levels were determined by qPCR. n=3-4/group, technical replicates (tech rep)=2/sample. **C)** Liver sections were prepared from 3 fasted mice per group and stained with Oil-red-O neutral lipid dye. Representative images are depicted. **D)** Liver triglyceride (TG) concentrations were determined biochemically. n=3-4/group, except that n=2 at t=6 and t=12 h, tech rep=2/sample. **E)** Plasma non-esterified fatty-acid (NEFA) levels were measured biochemically. n=3-5/group, tech rep=2/sample. **F)** Plasma NEFA composition of starved animals was determined by GC-FID. n=4/group, tech rep=1-2/sample. **G)** MM liver fractions from (A) were analyzed using p-AKT (S473), AKT, p-S6 (S240-S244) and S6 specific antibodies by western blot. Unpaired t-tests were used to compute significance levels, not significant, n.s.; p≤0.05 *; ≤0.01 **; ≤0.001 ***.

ATGL is not only important in adipose-tissue, but it is also a critical TG-hydrolase in the liver (Ong, Mashek, Bu, Greenberg, & Mashek, 2011; Zimmermann et al., 2004). To test if it plays a role in liver SREBP-1c regulation by catalyzing the release of uFAs from liver TG stores, we used ALKO mice, conditionally lacking *Atgl* only in the liver (*Atgl*^flox/flox, AlbuminiCre+/-^) and isogenic controls (*Atgl*^flox/flox^) (Wu et al., 2011). The animals were again subjected to a fasting / refeeding regimen and in liver WBs, P- and N-SREBP-1c signals increased in both genotypes after refeeding (Figure 4A). When comparing ALKO mice to controls, the P-SREBP-1c signal was elevated 6 h after refeeding, and the N-SREBP-1c signal was weakly elevated 6 h and 9 h after refeeding. In spite of this, the expression of the SREBP-1c target genes, *Acc-1* and *Fasn*, was not significantly different between the genotypes (Figure 4B). Nevertheless, the ALKOs were consistently slightly higher than the controls. ATGL deficient ALKO livers, appeared pale (Figure 4C, upper panel), and respective sections stained much stronger with ORO (Figure 4C, lower panel), as their TG content was consistently higher than in control livers (Figure 4D) (Ong et al., 2011; Wu et al., 2011). Since ATGL was lacking in livers (Figure 4A, lower panels) but not in adipose-tissue (Wu et al., 2011), ALKO mice showed similar plasma NEFA concentrations as controls (Figure 4E). WB analysis of liver-specific insulin receptor signaling revealed increased p-AKT signals 6 h after HChD refeeding in ALKO compared to control (Figure 4F, upper panels). Analysis of p-S6 showed no difference between the genotypes (Figure 4F, lower panels). These data suggest that in the presence of functional ATGL in adipose-tissue, lack of liver ATGL only leads to a very weak but still relatively persistent upregulation of N-SREBP-1c levels. Moreover, tissue-specific insulin signaling seems not to contribute to this residual ATGL effect.

**Figure 4:**
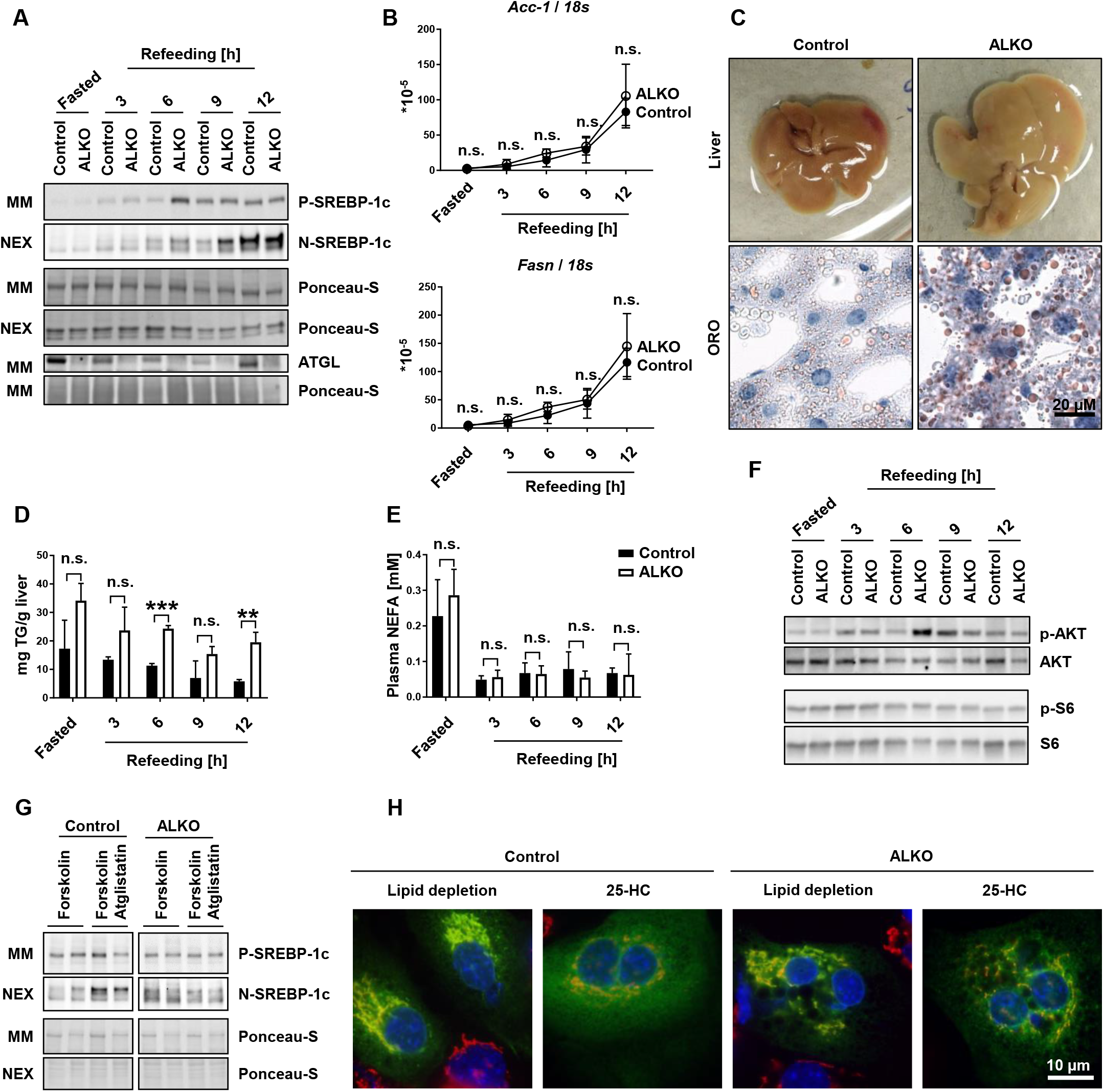
SREBP-1c processing in liver specific *Atgl* knockout mice and hepatocytes. Liver specific knockout of *Atgl* (ALKO) leads to a weak increase of N-SREBP-1c during high carbohydrate/low-fat diet (HChD) re-feeding, compared to controls. Mice were fasted for 12 h over-night and subsequently groups of mice were either sacrificed (Fasted) or, refed an HChD and sacrificed at the time-points indicated (Refeeding). **A)** Livers were resected and microsomal membrane fractions (MM) and soluble nuclear extracts (NEX) were prepared. Respective extracts from 3-5 mice were pooled and analyzed by western blot using antibodies specific for the proteins indicated. Ponceau-S stained membranes are shown as loading controls. **B)** Liver *Acc-1* and *Fasn* (mRNA) levels were determined by qPCR. n=3-4/group, technical replicates (tech rep)=2/sample. **C)** Liver sections from three fasted mice per genotype were stained with Oil-red-O neutral lipid dye (ORO). Representative liver- and ORO- images are depicted. **D)** Liver triglyceride (TG) concentrations were determined biochemically. n=4/group, tech rep=1/sample. **E)** Plasma non-esterified fatty-acid (NEFA) levels were measured biochemically. n=3-4/group, tech rep=1/sample. **F)** MM liver fraction pools from (A) were incubated with p-AKT (S473), AKT, p-S6 (S240-S244) and S6 specific antibodies by western blot. **G)** Primary hepatocytes were isolated and cultured under lipid sufficiency in hepatocyte medium. On the next day, cells were set up in 5% lipoprotein deficient serum (LPDS) medium with forskolin present to activate intracellular lipolysis and either treated with vehicle or the ATGL specific inhibitor Atglistatin. Cells were incubated in these media for 16 h and 2 h before harvest, the protease inhibitor ALLN was added. MM and NEX were prepared and subjected to western blot. Each lane corresponds to one culture dish. **H)** Primary hepatocytes were transfected with pGFP-SCAP one day after isolation. 48 h later cells were treated with 1% w/v (2-hydroxypropyl)-beta-cyclodextrin for 1 h. Next, cells were re-fed with 5% LPDS medium containing mevalonate and mevastatine (lipid depletion), and if indicated, 10 µg/ml 25-hydroxycholesterol (25-HC). 2 h later, cells were formaldehyde-fixed and permeabilized with Triton X-100. GFP-SCAP was visualized by immunofluorescence using anti-GFP antibodies followed by Alexa-488 coupled secondary antibodies (green); Golgi was imaged by anti-GM130 followed by Alexa-594 coupled secondary antibodies (red); DAPI was used for nuclear staining. Images of 20 cells/condition were analyzed. Representative images are shown. See Figure 4-figure supplement 1A, for all color channels, and Figure 4-figure supplement 1B for Pearson correlation analysis of co-stainings. Unpaired t-tests were used to compute significance levels, not significant, not significant, n.s.; p≤0.05 *; ≤0.01 **; ≤0.001 ***.

To analyze the importance of ATGL for SREBP-1c activation at the cellular level, we cultured primary hepatocytes from ALKO or control mice under lipid depleted conditions (5% LPDS) with forskolin present, to activate intracellular lipolysis. Treatment with selective ATGL inhibitor Atglistatin (Mayer et al., 2013; Schweiger et al., 2017) led to an increase in N-SREBP-1c WB signal in control cells but not in those lacking ATGL (Figure 4G). This result demonstrates that ATGL activity, presumably by producing uFAs, represses SREBP-1c activation in hepatocytes when they are decoupled from systemic regulation by adipose-tissue derived FA.

To follow intracellular SCAP trafficking, hepatocytes were transfected with pGFP-SCAP and rigorous lipid depletion was applied, first using (2-hydroxypropyl)-beta-cyclodextrin, and then 5% LPDS (Shao, Machamer, & Espenshade, 2016). Under these conditions, GFP-SCAP (green) co-localized with the Golgi marker GM130 (red) in control hepatocytes and to a somewhat higher extend in ALKO hepatocytes (Figure 4H and Figure 4-figure supplement 1A,B). Addition of 25-HC reduced GFP-SCAP/GM130 co-localization in control hepatocytes stronger than in ALKO hepatocytes (Figure 4H, and Figure 4-figure supplement 1A,B). In conclusion, SREBP-1c was activated in the presence of genetic or pharmacological ATGL inhibition and the essential activation step, transport of SCAP from ER to Golgi (Shao et al., 2016), was also controlled by ATGL.

## Discussion

Overall, our findings explain one important aspect of the interplay between lipolysis and lipogenesis: ATGL activity in adipose-tissue liberates uFAs that suppress SREBP-1c activation in the liver. As a result, in wild-type animals, the fatty-acid synthase machinery in the liver is suppressed as long as adipose-tissue-derived uFAs are abundant (see, *Fasn* and *Acc-1* qPCRs, Figures 2B and 3B). Additionally, we found that liver cell ATGL activity can also mildly suppress liver SREBP-1c cleavage activation *in vivo* (Figure 4A) and in primary hepatocytes (Figure 4G). Our studies of GFP-SCAP trafficking indicate that lipolysis-derived uFA act through control of SCAP-SREBP ER-to-Golgi transport (Figure 4H, and Figure 4H and Figure 4-figure supplement 1A, B) (J. D. Horton et al., 2002). Published *in vitro* studies demonstrate that uFAs regulate SCAP transport by controlling the stability of INSIG-1 through a mechanism that requires the ER-associated degradation machinery protein UBXD-8 (Lee et al., 2010; Lee et al., 2006). Importantly, the tissue source of these SREBP cleavage suppressing uFAs was previously unknown. Our studies fill this gap and demonstrate *in vivo* the importance of adipose-derived uFAs and ATGL for homeostatic control of hepatic lipogenesis.

## Materials and Methods

| Reagent type | Designation | Host organism | Source | Catalog number | RRID | Additional information |
| --- | --- | --- | --- | --- | --- | --- |
| Antibody | Anti-SREBP-1c, clone 2A4 | mouse | Abcam | ab3259 | AB_303650 | 1:500 |
| Antibody | Anti-SREBP-1c clone 20B12 | rabbit | Merck | MABS1987 | AB_2877057 | 1:500 |
| Antibody | Anti-FLAG® M2 | mouse | Sigma-Aldrich | F3165 | AB_259529 | 1:500 |
| Antibody | Anti-S6 ribosomal protein, clone 5G10 | rabbit | Cell Signaling | 2217 | AB_331355 | 1:1000 |
| Antibody | Anti-phospho-S6 Ribosomal Protein, (Ser240/244) (D68F8) | rabbit | Cell Signaling | 5364 | AB_10694233 | 1:5000 |
| Antibody | Anti-AKT | rabbit | Cell Signaling | 9272 | AB_329827 | 1:1000 |
| Antibody | Anti-phospho-Akt, (Ser473), clone D9E | rabbit | Cell Signaling | 4060 | AB_2315049 | 1:1000 |
| Antibody | Anti-GFP-avictoria antibody | rabbit | Abcam | ab290 | AB_303395 | 1:500 |
| Antibody | Anti-GM130, clone 35 | mouse | BD Biosciences | 610822 | AB_398141 | 1:250 |
| Antibody | Anti-Rabbit IgG (H+L), Alexa Fluor 488 coupled | goat | Invitrogen | A11034 | AB_2576217 | 1:250 |
| Antibody | Anti-Mouse IgG (H+L), Alexa Fluor 594 coupled | mouse | Invitrogen | A11005 | AB_141372 | 1:250 |
| Antibody | Anti-mouse Immunoglobulins/ HRP | goat | Dako | P0477 | AB_2617137 | 1:3000 |
| Antibody | Anti-rabbit Immunoglobulins/ HRP | swine | Dako | P0217 | AB_2728719 | 1:3000 |
| Cell line | U2OS |  | ATCC | HTB96 | CVCL_0042 |  |
| Mouse line | AKO/cTg |  | Rudolf Zechner | MGI: 5294406 | <u>MGI:5294406</u> | *Atgl* knockout mice with transgenic *Atgl* expression in the heart |
| Mouse line | AAKO |  | Rudolf Zechner | JAX: 024278<br>x<br>JAX: 028020 | <u>IMSR JAX:024278</u><br>x<br><u>IMSR JAX:028020</u> | Adipose specific *Atgl* knockout mice; own breeding |
| Mouse line | ALKO |  | Rudolf Zechner | JAX: 024278<br>x<br>JAX: 018961 | <u>IMSR JAX:024278</u><br>x<br><u>IMSR JAX:018961</u> | Liver specific *Atgl* knockout mice; own breeding |

### Chemicals

We obtained mevastatin (M2537), mevalonate (50838) and *N*-acetyl-leucinyl-leucinyl-norleucinal (ALLN, 208719) from Merck, Germany; 25-hydroxycholesterol (25-HC, H1015), lipoprotein-deficient serum (LPDS; S5394), ITS Liquid Media Supplement (100×) (ITS, I3146), Forskolin (F3917) and all other powdered chemical substances from Sigma Aldrich, USA. Atglistatin was synthesised by the group of Dr. Breinbauer, Graz University of Technology, Austria.

### Cell culture media and supplements

Fetal bovine serum (FBS, 10500), high glucose Dulbecco’s modified Eagle’s medium (DMEM, 41966052) and penicillin-streptomycin (PenStrep, 15140-122) were from Gibco, USA. Standard cell culture medium (D10F) contained, DMEM containing 1x PenStrep and supplemented with 10% (v/v) FBS. Hepatocyte medium: DMEM containing 1x PenStrep, 20% (v/v) FBS, 100 nM dexamethason, 1x ITS-Supplement. 5% LPDS: DMEM containing 1x PenStrep and supplemented with 5% (v/v) LPDS (Sigma Aldrich, USA, S5394).

### Stable 2xFlag-SREBP-1c U2OS cell line generation

We used the human SREBP-1c cDNA containing vector pQCXIN (Addgene, USA, 631514) as a template to generate 2x Flag tagged, full length, human *SREBP-1c* by conventional PCR, using 2xFlag-h*SBP-1c*-FW and hSBP-1c-REV primers, listed below. The PCR product was introduced into pCDNA3.1 (Invitrogen, USA, V790-20) using the NEB, USA, builder® HiFi DNA Assembly Cloning Kit (NEB, USA, E5520S). The resulting pCDNA3.1 2x Flag-*SREBP-1c* was stably transfected into U2OS cells (ATCC®, USA, HTB96™) using Lipofectamine 2000 (Thermo Fisher Scientific, USA, 11668019) and selected for resistance to G 418 disulfate salt (Sigma Aldrich, USA, A1720). Finally, we generated the UF1c cell-line by clonal selection for 2x Flag-SREBP-1c expression. 2xFlag-h*SBP-1c*-FW: AAGCTTGGTACCGAGCTCGCACCATGGATTATAAAGATCATGATATCGATTACAAGGATGACGATGACAA; h*SBP-1c* REV: CGGCCGCCACTGTGCTGGATCTAGCTGGAAGTGACAGTGG.

### Uf1c cell line maintenance and experiments

Uf1c cells were cultured under standard cell culture conditions in a humidified chamber at 37°C, 5% CO2 in D10F medium. For experiments, cells were seeded at 60% confluency in 6-well plates. On the next day, cells were washed twice with PBS and subsequently incubated in 5% LPDS for 16 h, containing 100 µM BSA bound FA or, 1 mg/ml 25-HC, if mentioned in the Figure. 2 h before harvest, 25 µg/ml ALLN was added. Cells were harvested directly in FSB (final sample buffer: 60 mM Tris-HCl at pH 7,4; 2% (w/v) SDS; 10% glycerol) and subsequently, analysed by WB.

### BSA bound FA

4 mM FA sodium salt (sodium palmitate (Cayman, P9767); palmitoleic acid sodium salt (Sigma Aldrich, USA, 6610-24-8); sodium oleate (Sigma Aldrich, USA, O75011); linoleic acid sodium salt (Sigma Aldrich, USA, L8134)) solutions in double distilled H2O, were mixed with 172 mg/ml (w/v) FA free BSA (Sigma Aldrich, USA, A7030) in 2x PBS at 37° under constant vortexing, to achieve BSA bound FA in PBS. FA concentrations were determined using the NEFA kit HR Series NEFA-HR(2) from (WAKO Chemicals, Japan, 276-76491, 995-34791, 993-35191, 999-34691, 991-3489)

### Ethical approval

All animal studies were performed in accordance with the guidelines and provisions of the Commission for Animal Experiments of the Austrian Ministry of Education, Science and Research (BMBWF). Approved animal applications and amendments include, BMBWF-66.007/0015-V/3b/2018; BMBWF-66.007/0004-V/3b/2019 and BMBWF-2020-0380.481.

### Animal experiments

Mice were either fed (non-fasted), fasted or, fasted and subsequently refed (refed). The non-fasted group was fed ad libitum with a standard chow diet ((4.5% fat, 34% starch, 5.0% sugar, and 22.0% protein) M-Z extrudate, V1126, Ssniff Spezialdiäten, Germany), the fasted group was fasted for 12 h, from 7 a.m. to 7 p.m. The refed group was fasted for 12 h from 7 a.m. to 7 p.m. and then refed an high-carbohydrate/low-fat diet (HChD, equivalent to TD 88122; Harlan Teklad, USA) up to 12 h. Mouse strains used: WT: C57Bl/6J (own breeding, originally from Jackson lab). Genetically modified strains on C57Bl/6J background: AKO/cTg, *Atgl*-knockout/cardiac transgenic *Atgl* mice (*Atgl*^-/-,Myh6ATGL+/^) (Haemmerle et al., 2011; Schreiber et al., 2015b); AAKO, Adipose-tissue specific *Atgl*-knockout (*Atgl*^flox/flox,AdipoQCre+/-^) (Schoiswohl et al., 2015); ALKO, Liver specific *Atgl*-knockout (*Atgl*^flox/flox, AlbuminiCre^) (Wu et al., 2011). All mice were between 8 and 22 weeks of age. Group allocation was based on genotype, solely. Apart from that, mice were chosen randomly for experiments.**Hepatocyte isolation** Primary hepatocytes were isolated by perfusion of mouse livers with 40 ml perfusion buffer (5.5 mM KCl, 0.1% Glucose, 2.1 g/l NaHCO3, 700 μM EDTA, 10 mM Hepes and 150 mM NaCl). After 20 min the buffer was exchanged to 50 ml collagenase buffer (5.5 mM KCl, 0.1% Glucose, 2.1 g/l NaHCO3, 10 mM Hepes and 150 mM NaCl, 3.5 mM CaCl2, 1% BSA, 500 μg/ml Collagenase Type I (300 U/mg)). Livers were perfused at a perfusion rate of 2 ml/min. Subsequently, livers were dissociated with a plunger of a syringe in 10 ml D10F. The homogenate was applied onto a 100 μM cell strainer and the flow through collected. Hepatocytes were centrifuged at 100 *g* for 2 min and washed twice in DMEM containing 1x PenStrep. Next, cells were re-suspended in hepatocyte medium. Cells were counted after staining with trypan blue (Thermo Fischer Scientific, USA, 15250061) to assess viability. Primary hepatocytes were seeded in rat tail collagen I (Sigma Aldrich, USA, C3867) coated 6-well dishes at a density of 6*10^5^ cells/well.

### Hepatocyte Atglistatin experiment

One day after isolation, hepatocytes where washed twice with PBS and cultured in 5% LPDS medium containing 10 µM Forskolin or 10 µM Forskolin and 50 µM Atglistatin. After 14 h, 25 µg/ml ALLN was added for two hours. Then microsomal (MM) and nuclear fractions (NEX) were isolated.

### Microsomal- and nuclear- fractionation of mouse hepatocytes

Microsomal membrane fractions and nuclear fractions were isolated from hepatocytes, essentially, as described by Hannah *et al*. (Hannah et al., 2001). In brief, cells were washed with buffer B (10 mM Hepes-KOH pH 7.4, 10 mM KCl, 1.5 mM MgCl2, 0.5 mM EDTA, 0.5 mM EGTA, 1 mM Dithiothreitol (DTT), 25 µg/ml ALLN and protease inhibitor cocktail present (Thermo Fisher Scientific, USA, A32963)). Next, cells were incubated 10 min on ice in buffer B. The cell suspension was passed through a 23 gauge needle using a 1 ml syringe 7 times and centrifuged at 1000 *g* at 4°C for 5 min. The resulting 1,000 *g* pellet was re-suspended in buffer C (10 mM Hepes-KOH at pH 7.4, 0.42 M NaCl, 2.5% (v/v) glycerol, 1.5 mM MgCl2, 0.5 mM sodium EDTA, 0.5 mM EGTA, 1 mM DTT and the protease inhibitor cocktail (Thermo FisherScientific, USA, A32963)), and slowly rotated at 4°C for 45 min, followed by centrifugation at top speed (17,000 g) in a table top micro centrifuge at 4°C for 30 min. The resulting 17,000 *g* supernatant was designated soluble nuclear extract, and made 1 x FSB. The former 1,000 *g* supernatant was used to prepare the microsomal membrane fraction by centrifugation at 10^5^ *g* in a Beckman TLA 100.4 rotor for 30 min at 4°C, followed by resuspension of the 10^5^ *g* pellet in FSB.

### Immunofluorescence (IF)

IF was performed and, co-localization of GFP-SCAP and GM130 analysed, essentially as previously described by Shao *et al*., [4], with modifications as follows. 2,5*10^5^ hepatocytes were seeded into 1 well of a 6 well-plate containing a sterile coverslip using standard hepatocyte medium. Hepatocytes were transfected one day after isolation using Lipofectamine 2000 (Thermo Fisher, USA, 11668019) with 1 µg pGFP-SCAP plasmid. 48 h post transfection cells were treated with 1% hydroxypropyl-beta-cyclodextrin (HPCD) to deplete sterols for 1 h in plain DMEM medium. Next, cells were re-fed for 2 h with 5% LPDS medium supplemented with 50 µM mevalonate and 50 µM mevastatin. Furthermore, 10 mg/ml 25-HC was added when indicated. Thereafter, cells were washed with PBS twice, fixed in formaldehyde/PBS (0.03/1; w/v) at room temperature for 10 min and then permeabilized with Triton X-100/PBS/glycine (0.05/0.9/0.1; v/v/v) for 3 min at room temperature. Next, cells were incubated for 30 min with primary antibodies (anti-GFP and anti-GM130) and respective (Alexa-488 (green) and Alexa-594 (red) coupled) secondary antibodies, followed by DAPI (Sigma Aldrich, USA, D9542) staining. For details, see reagent table, above. Coverslips were mounted onto glass-slides and dried in the dark over-night before visualization by an Olympus BX51 microscope at the microscopy core-facilty of the ZMF, MUG, Graz, Austria. Quantitative co-localization analysis was conducted using 20 pictures of each condition. The analysis was performed using Image J with the JACoP plug-in (Shao et al., 2016).

### Liver microsomal- and nuclear- fractionations

Liver cell fractions were prepared as previously described, with small modifications (Engelking et al., 2004). Livers were excised and washed in ice cold PBS and immediately frozen in liquid nitrogen cooled methylbutane. To isolate nuclear fractions, 1 g of frozen liver was mixed with 6 ml buffer A (10 mM Hepes at pH 7.6, 25 mM KCl, 1 mM sodium EDTA, 2 M sucrose, 10% vol/vol glycerol, 0.15 mM spermine, 2 mM spermidine, 1x protease Inhibitor cocktail, 50 µg/ml ALLN). Livers were homogenized by three strokes with a teflon pestle in a potter homogenizer at low speed. The homogenate was filtered through a 100 µM cell strainer. Samples were overlayed with 2 ml buffer A in SW 41Ti tubes and the tubes filled with Buffer 1 (10 mM Hepes at pH 7.6, 25 mM KCl, 1 mM sodium EDTA). The samples were then centrifuged at 25,000 rpm (75,000 g) for 1 h at 4°C in a SW 41 Ti Rotor. Subsequently, the tubes were turned over, the nuclear pellet recovered from the bottom and resuspended in 1 ml buffer D (10 mM Hepes pH 7.6, 100 mM KCl, 2 mM MgCl2, 1 mM sodium EDTA, 10% (vol/vol) glycerol, 1 mM DTT, 1x protease inhibitor cocktail, 50 µg/ml ALLN). For extraction of soluble nuclear proteins, 140 µl buffer AS (3.3 M ammonium sulfate (pH 7.9) was added to the solution, agitated gently for 40 min at 4°C on a rotating wheel in cold room and subsequently centrifuged at 78,000 rpm in Beckman TLA-100.4 rotor for 45 min at 4°C. The supernatant was mixed 1:5 with 5x FSB and designated soluble nuclear protein extract (NEX). For membrane fractions 50 mg frozen livers were homogenized with three potter strokes in 1 ml buffer M (20 mM Tris-HCl pH 7.4, 2 mM MgCl2, 0.25 mM sucrose, 10 mM sodium EDTA, 10 mM sodium EGTA, 1x protease inhibitor cocktail, 50 µg/ml ALLN). The homogenate was centrifuged at 6,800 *g* for 5 min at 4°C in a micro centrifuge. The resulting supernatant was centrifuged at 48,600 rpm for 30 min at 4°C in the Beckman TLA-100.4 rotor. The pellet was dissolved in 1x FSB and designated cytosolic microsomal membrane extract (MM).

### Blood Biochemistry

Plasma NEFA levels were analysed using the NEFA kit HR Series NEFA-HR(2) from (276-76491, 995-34791, 993-35191, 999-34691, 991-34891, WAKO Chemicals, Japan) according to manufactures instructions. Liver TG levels were measured from liver Folch extracts (Folch, Lees, & Sloane Stanley, 1957) using the Triglycerides FS 10 kit (Diasys, Germany, 15760991002). Samples where dissolved in 600 µl 1% Triton X-100. Plasma insulin levels were determined using the mouse insulin ELISA kit (Chrystal Chem, USA, 90080).

### Fatty-acid analysis

Livers were homogenized, subjected to Folch extraction and lipids dried under a stream of nitrogen. Lipid extracts were pre-separated by thin layer chromatography (TLC), The band co-migrating with a triolein standard was scraped off and after addition of C15:0 as internal standard directly trans-esterified (1.2 ml toluene and 1 ml boron trifluoride-methanol (20%)) at 110 °C for 1 h. GC analysis of the corresponding fatty-acid (FA) methyl esters was performed as described (8) and concentrations were quantitated by peak area comparison with the internal standard.

### Western Blot (WB)

WB was performed using 4-20% SDS gels (Bio-Rad, 4561096) and blotted on 0.45 µm nitrocellulose membranes (GE healthcare, 15259794). Next, membranes were stained with Ponceau-S solution (Sigma Aldrich, USA, P7170) for protein load normalization and, subsequently blocked using skim milk powder (Sigma Aldrich, USA, 1153630500) /PBS/TWEEN^®^ 20 (Sigma Aldrich, USA, P1379) (0.05,1,0.05; w/v/v). Membranes were incubated with one of the following first antibodies overnight: Anti- N-terminal SREBP-1c, anti-Flag-Tag M2, anti-S6 ribosomal protein, anti-phospho-S6 ribosomal protein (P-S6) S240/244, anti-AKT and anti-phospho-AKT (P-AKT) S473. As secondary antibodies, respective horseradish peroxidase coupled secondary antibodies were used. Signals were detected using the ChemiDoc Imaging System (Biorad, USA, 17001401). WB band intensities were analysed using image J, NIH, software package (Schneider, Rasband, & Eliceiri, 2012). Relative levels of N-SREBP-1c were determined by division of N-SREBP-1c band intensities through respective P-SREBP-1c band intensities, essentially as described earlier (Hannah et al., 2001). Stripped (Thermo Scientific, USA, 46430) membranes were used for phosphorylated protein blots, except for Figure 4, AKO/cTg S6 and P-S6 blots, two different membranes with similar loading were used.

### Quantitative Polymerase Chain Reaction (qPCR)

RNA was isolated from livers homogenized in Buffer M using Trizol (Invitrogen, USA, 15239794), cDNA prepared using High-Capacity cDNA Reverse Transcription Kit (Applied Biosystems, USA, 4368814) and qPCR performed using the SYBR Green Luna® Universal qPCR Master Mix (NEB, USA, M3003) on the QuantStudio™ 7 Flex Real-Time PCR System (Applied Biosystems™, USA, 4485701). Primers were designed with the NCBI primer designing tool (primer-blast) and are listed in table 1. Relative mRNA levels were analyzed as described by Schmittgen *et al*.,(Schmittgen & Livak, 2008) with modifications, as follows. The relative PCR-efficiency of the gene of interest (GOI) primer sets compared to the 18s rRNA (HKG) primer set was determined by computational standard curve analysis in QuantStudio™ 7 Flex Software, Applied Biosystems™, USA) and subsequent division of GOI/HKG primer efficiencies. The resulting value was used as a base to compute relative quantities from delta CT values.

**Table 1.**
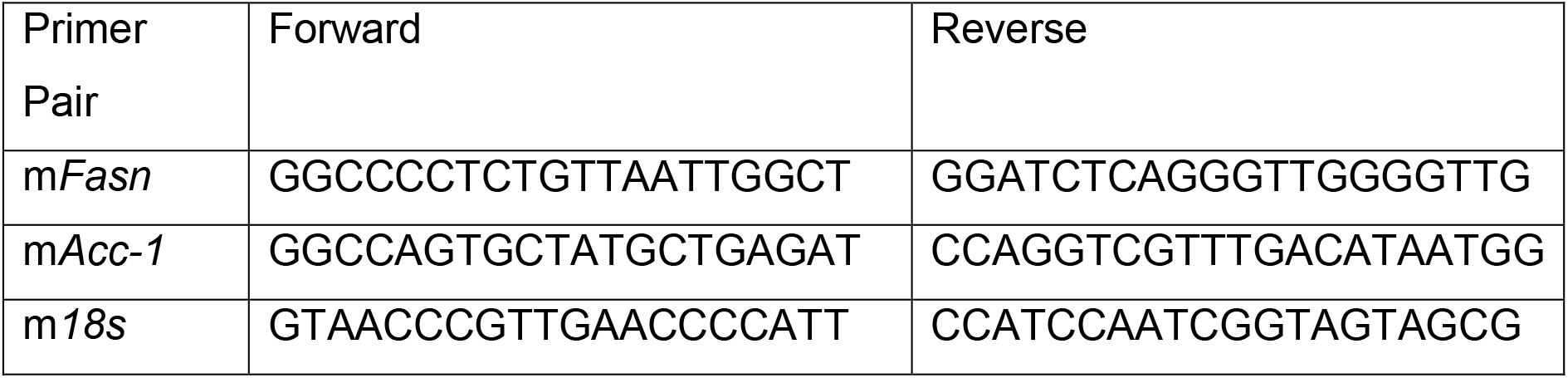
qPCR Primer sequences.

### Oil-Red-O staining

Oil-Red-O (ORO-staining) (Sigma-Aldrich, USA, O0625) was performed on 4% neutral buffered formalin fixed, frozen sections from mouse livers that were fresh frozen in lN2 cooled methylbuthane, using a standard protocol (Schreiber et al., 2015a).

### Statistical analyses

A biological replicate is defined as a biological unit, *i.e*. one cell or one liver or one mouse. Biological replicate values were computed as the arithmetic mean value of technical replicate values. Mean values of biological replicate values were calculated using the “graph pad prism 7.04” function mean with SD (standard deviation) and the number of biological replicate values underlying each mean value is given as “n=x”. A technical replicate value however, represents the result of one (of many) measurement(s) taken from one and the same biological unit. The number of technical replicate values underlying each biological replicate value is given as “technical replicates (tech rep) = x”. Biological outliers were detected using the Graph Pad online tool at “https://www.graphpad.com/quickcalcs/grubbs1/” that is based on the “Grubbs’ test, also called the ESD method”. The significance level chosen was, Alpha = 0.05 (standard).

## Acknowledgments

The authors would like to thank Jared Rutter for kindly providing *SREBP-1c* cDNA; Wei Shao, Chiaki Ishida, Chune Liu, Debaditya Mukhopadhyay and Shan Zhao for their continued assistance and advice during experiments done in Dr. Espenshade’s laboratory; Peter Hofer, Guenter Haemmerle and Kathrin Zierler for technical assistance and critical advice; Markus Absenger for technical assistance and Anna Migglautsch for Atglistatin synthesis. This work was funded by the Austrian Science Fund (FWF); DK-MCD W1226 and the Medical University of Graz. Beatrix Wieser received additional funding through the Austrian Marshall Plan Scholarship Program (953 1175 38 22 2019) and the Bundesministerium fuer Bildung, Wissenschaft und Forschung (BMBWF), Oesterreichische Austausch Dienst-GmbH (OeAD-GmbH), Zentrum fuer Internationale Kooperation und Mobilitaet (ICM) through a Marietta Blau Grant (ICM-2019-13518). Paul Willibald Vesely was supported by the European Research Council Grant, LipoCheX (340896), granted to Rudolf Zechner, and is supported by the FWF Grant, LipoLung (P30968), granted to Gerald Hoefler.

## Author contributions

B.I.W.; P.W.V; G.H. M.S.; R.Z.; R.P.; P.J.E. contributed to experimental design, data interpretation and manuscript preparation.

B.I.W.; P.W.V. and S.S. performed and analysed qPCR data.

B.I.W. and P.P.S performed cell culture experiments and WBs

In vivo studies were conducted by B.I.W.; P.W.V. and S.S.

Immunofluorescence was performed by S.S.; B.I.W. and P.W.V.

Hepatocyte experiments where performed by B.I.W.

Blood samples were analysed biochemically by B.I.W.

ORO-staining was performed by S.S.

FID/GC was performed by W.S and H.R.

Statistical analysis was performed by B.I.W. and P.W.V.

