## Supplementary material for "Adipose Triglyceride Lipase is needed for homeostatic control of Sterol Element-Binding Protein-1c driven hepatic lipogenesis": Figure 4 Supplement 1.

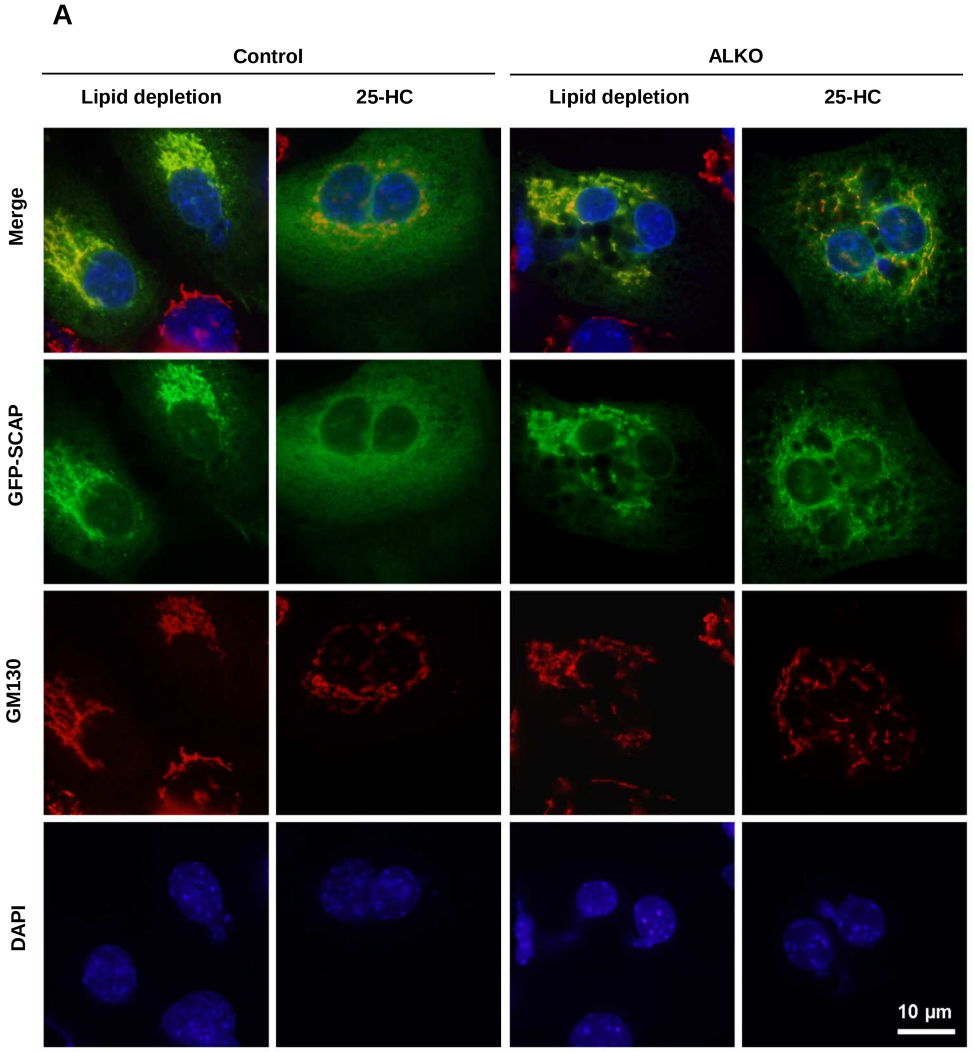


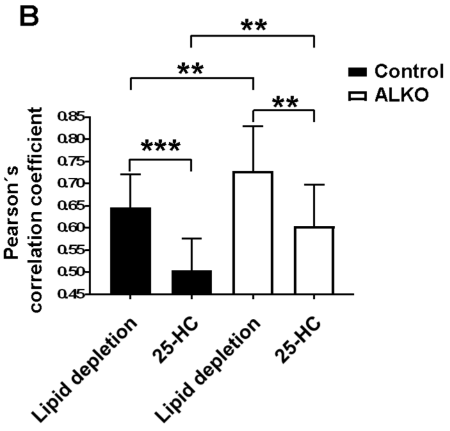


**Figure 4-figure supplement 1: GFP-SCAP/GM130 co-localization in primary hepatocytes. (A)** Primary hepatocytes were transfected with pGFP-SCAP one day after isolation. 48 h later cells were treated with 1% w/v (2-hydroxypropyl)-beta-cyclodextrin for 1 h. Next, cells were re-fed with 5% LPDS medium containing mevalonate and mevastatine (lipid depletion), and if indicated, 10 µg/ml 25-hydroxycholesterol (25-HC). 2 h later, cells were formaldehyde-fixed and permeabilized with Triton X-100. GFP-SCAP was visualized by immunofluorescence using anti-GFP antibodies followed by Alexa-488 coupled secondary antibodies (green); Golgi was imaged by anti-GM130 followed by Alexa-594 coupled secondary antibodies (red); DAPI was used for nuclear staining. Images of 20 cells/condition were analyzed. Representative images are shown. **(B)** GFP-SCAP/GM130 co-localization was quantified with the Pearson´s correlation coeffiecent using ImageJ. n=20 cells/group, technical replicates=1/cell. Unpaired t-tests were used to compute significance levels, not significant, n.s.; $p \leq 0.05$ *; $\leq 0.01$ **; $\leq 0.001$ ***.
